# SARS-CoV-2 predation of Golgi-bound PI4P primes the massive activation of the DNA Damage Response kinase ATM in the cytoplasm

**DOI:** 10.1101/2024.12.05.626967

**Authors:** Antoine Rebendenne, Caroline Soulet, Ana-Luiza Chaves Valadaõ, Boris Bonaventure, Joe McKellar, Olivier Moncorgé, Caroline Goujon, María Moriel-Carretero

**Affiliations:** IRIM, Université de Montpellier, CNRS; CRBM, Université de Montpellier, CNRS

## Abstract

Like all viruses, SARS-CoV-2, the causative agent of COVID-19, relies on host cell resources to replicate. Our study reveals that, among these resources, SARS-CoV-2 hijacks the oxysterol-binding protein 1 (OSBP1) transporter to exploit the Golgi-bound phosphatidylinositol-4-phosphate (PI4P) pool. This leads to a depletion of Golgi-resident PI4P, triggering the activation of the ATM DNA Damage Response (DDR) kinase in the cytoplasm. As such, ATM, typically anchored to PI4P at the Golgi in an inactive state, undergoes auto-phosphorylation and cytoplasmic release upon SARS-CoV-2-induced PI4P depletion. Conversely, pharmacological inhibition of ATM auto-phosphorylation, which stabilizes its interaction with PI4P, significantly impairs SARS-CoV-2 replication. The requirement for PI4P and impact of ATM inhibition might be conserved across coronaviruses, as similar effects were observed with HCoV-229E. Finally, SARS-CoV-2-induced, cytoplasmic ATM pre-activation primes cells for an accelerated response to DNA damage, which might contribute to the severe outcomes of COVID-19 observed in cancer patients undergoing chemo- or radiotherapy. Therefore, this study uncovers a DNA damage-independent mode of ATM activation and highlights the potential of ATM inhibitors as therapeutic agents against COVID-19.

## Introduction

The severe acute respiratory syndrome coronavirus 2 (SARS-CoV-2) is the causative agent of the global COVID-19 pandemic. After SARS-CoV-1, which caused an outbreak in 2002-2003, and Middle East Respiratory Syndrome coronavirus (MERS-CoV), discovered in 2012, SARS-CoV-2 has been the third highly pathogenic *Betacoronavirus* to cross the species barrier and emerge in the human population over the past 2 decades. Four other human coronaviruses (HCoVs) circulate in humans and cause common colds in winter: the *Alphacoronaviruses* HCoV-229E, and -NL63, and the *Betacoronaviruses* HCoV-OC43, and - HKU1. Coronaviruses enter their target cells after recognition of specific receptors (e.g. angiotensin converting enzyme 2 (ACE2), for SARS-CoV-1, SARS-CoV-2 and HCoV-NL63, or human aminopeptidase N (ANPEP) for HCoV-229E) by Spike envelope proteins. Fusion between the viral envelope and cellular membranes occurs either at the plasma membrane (when transmembrane protease serine 2, TMPRSS2, is present) or within the endosomes, and leads to the cytosolic delivery of the incoming genomic material. There, this positive, single stranded RNA is directly translated into pp1a and pp1ab polyproteins, which are processed into the individual non-structural proteins (nsps) forming the viral replication and transcription complex. In parallel, nsps hijack host cell membranes and components to induce the biogenesis of viral replication organelles constituted of perinuclear double membrane vesicles (DMVs). DMVs offer a protective environment for viral genomic RNA replication and transcription of subgenomic mRNAs (sg mRNAs) (reviewed in (Roingeard *et al*, 2022)). Subgenomic mRNAs are then notably translated into the structural proteins, which traffic through the endoplasmic reticulum (ER) membranes and the ER-to-Golgi intermediate compartment. In the latter, they associate with newly produced genomic RNA through interactions with the Nucleoprotein (N). This results in viral budding into the lumen of secretory vesicular compartments, and virions are finally secreted from the infected cell by exocytosis (reviewed in (V’kovski *et al*, 2021)).

To achieve their replication cycle, viruses heavily rely on host factors and components, including lipids. In particular, DMV-forming viruses induce host lipid synthesis and redistribution from different organelles in order to provide the lipids necessary for the major membrane rearrangements they require (reviewed in (Roingeard *et al*, 2022)). DMVs interact with host cell organelles at membrane contact sites (MCSs). Specific protein complexes, such as lipid transport proteins, are involved in managing these MCSs by acting on the apposed membranes. For instance, the oxysterol-binding protein 1 (OSBP1) transports newly synthesized cholesterol from the ER to the Golgi apparatus while transferring the phospholipid phosphatidylinositol 4-phosphate (PI4P) in the opposite direction. This mechanism is exploited by various single-stranded, positive-sense RNA viruses, such as some picornaviruses (the enteroviral genus members poliovirus, Coxsackievirus B3 -CVB3- and rhinovirus) or the flavivirus hepatitis C virus (HCV), which recruit PI4-kinase (PI4K) to their DMVs, leading to increased PI4P and enhanced OSBP1 recruitment for cholesterol transport. Hence, the coxsackievirus hijacks PI4KIIIβ (one of the four human PI4K isoforms) for this purpose. Similarly, HCV NS5A protein activates PI4KIIIα to elevate PI4P levels in DMVs, and the structural integrity of HCV DMVs is compromised when OSBP1 and PI4K are depleted, highlighting their role in maintaining DMV characteristics (Reiss *et al*, 2011).

The integrity of the cellular genetic material is ensured by surveillance and repair strategies capable of reacting to DNA damage or to situations likely to compromise DNA integrity. Such surveillance mechanism is collectively termed the DNA Damage Response (DDR) (Reinhardt & Yaffe, 2013). The damaged DNA is detected by dedicated sensors, for example the MRE11-RAD50-NBS1 (MRN) complex recognizing DNA double strand breaks, while exposed single-stranded DNA is rapidly coated by Replication Protein A. These in turn engage upstream orchestrating kinases, namely ATM and ATR, respectively. These kinases both possess auto-phosphorylation abilities, capable of boosting their own activation, and can also phosphorylate downstream effectors, such as CHK2 (for ATM) and CHK1 (for ATR) (Ciccia & Elledge, 2010). These and many other phosphorylated targets (Ciccia & Elledge, 2010) will engage transient cell cycle halting to give time for DNA repair, modulate transcriptional profiles and a myriad of other changes aimed at coping with the alert situation. Successful DNA repair will be followed by subsequent checkpoint inactivation, a situation termed recovery. However, persistence of the insult, or cellular defects in the repair machinery, create a situation in which damage cannot be repaired, frequently resulting in a permanent cell-cycle arrest that compromises cell homeostasis and can end up in cell death, or senescence, and inflammation, among others (Tomashevski *et al*, 2010; Kruman *et al*, 2004; Zhao *et al*, 2023b). As an alternative, cells can manage to resume cycling in the presence of the damage by illegitimate extinction of the DDR, an event called adaptation. Adaptation is typical of dysregulated scenarios, such as in cancer, and permits cell survival while propagating genome instability (Halazonetis *et al*, 2008).

Whereas the interplay between DNA viruses and the DDR has long been studied (Studstill *et al*, 2023), much less is known for RNA viruses. However, there is mounting evidence that RNA viruses can cause host genome instability and activation of diverse DDR branches, despite the fact that most of them do replicate in the cytoplasm (reviewed in (Ryan *et al*, 2016)). In particular, SARS-CoV-2 was shown to interact with components of the DDR (Grand, 2023). A comprehensive study using several human cellular models, as well as a mouse model and patient samples, showed that SARS-CoV-2 leads to the downregulation of the key DDR effector CHK1, rendering infected cells more prone to DNA damage (Gioia *et al*, 2023). Indeed, DNA damage was observed both in infected mice and in nasal and lung samples from COVID-19 patients (Gioia *et al*, 2023).

We recently discovered an unprecedented link between the membrane-anchored phospholipid PI4P and the master DDR kinase ATM (Ovejero *et al*, 2023). We found that, when not engaged in DNA damage signalling within the cell’s nucleus, ATM binds PI4P at the Golgi, where it lies in a resting state. Physiological, pharmacological and pathological situations that lead to a decrease in Golgi membrane-inserted PI4P release ATM, thus authorizing its auto-phosphorylation and making it more reactive to nuclear DNA damage (Ovejero *et al*, 2023). In view of the connection between some positive strand RNA viruses and PI4P, and the poorly characterized crosstalk of SARS-CoV-2 infection with the DDR, we investigated whether our knowledge of the PI4P-ATM interaction could prove useful to gain new knowledge about this infection. We discover that SARS-CoV-2 diverts PI4P available at the Trans-Golgi network by exploiting the OSBP1 transporter. As a consequence, PI4P is less available at the Golgi for its natural binders, including ATM, which is released, undergoes auto-phosphorylation and becomes massively enriched in the host cell’s cytoplasm. Reciprocally, inhibiting ATM’s auto-phosphorylation can stabilize the ATM-PI4P couple, and consequently antagonizes SARS-CoV-2 infection. We show that these concepts hold true in multiple cell types and apply to at least another coronavirus, HCoV-229E. Furthermore, we uncover that this mode of DNA damage-independent ATM activation can prime its reactivity to actual DNA damage, providing a potential rationale for the severity of the COVID-19 manifested by cancer patients undergoing chemo- and radiotherapy.

## Results

### SARS-CoV-2 exploits Golgi-derived PI4P

Positive strand RNA viruses forming DMVs, such as HCV, CVB3 or rhinoviruses, depend on PI4P for their replication (Hsu *et al*, 2010; Roulin *et al*, 2014). However, to our knowledge, whether SARS-CoV-2 relies on this molecule to replicate has not been assessed. To address this question, PI4P synthesis was inhibited using the drug PIK93 (Roulin *et al*, 2014) prior to infection of A549-ACE2 cells with a mNeonGreen (mNG)-expressing SARS-CoV-2 reporter virus, at multiplicities of infection (MOI) of 0.02 and 0.2. In the presence of PIK93, infection efficiency was significantly decreased as compared to the DMSO control (Figure 1A), suggesting a requirement for PI4P in SARS-CoV-2 replication.

**Figure 1.**
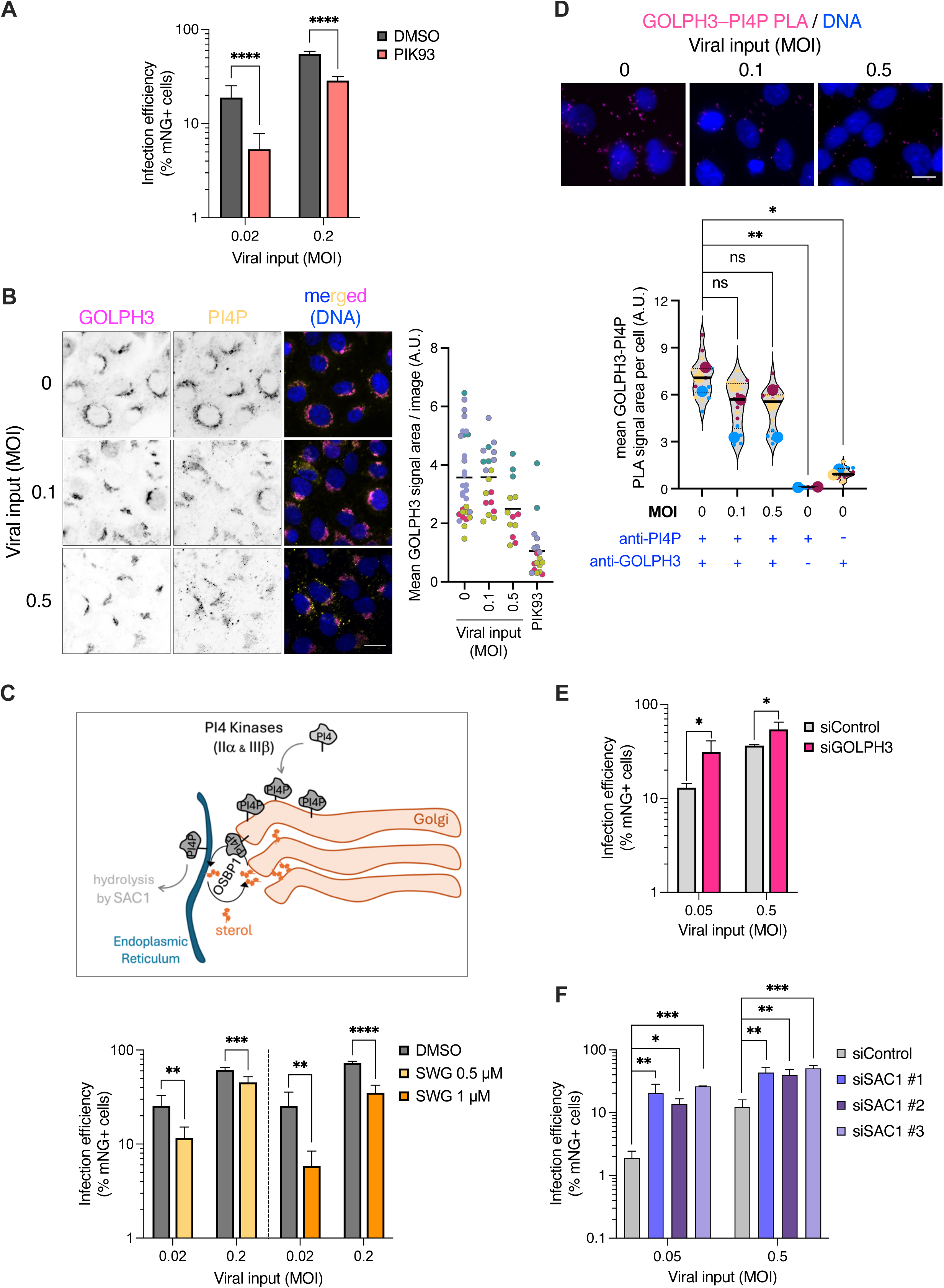
SARS-CoV-2 requires Golgi-bound PI4P to replicate. **A.** A549-ACE2 cells were incubated either with DMSO or 10 μM of PI4 kinase inhibitor PIK93 for 1 h prior to infection with SARS-CoV-2 bearing a mNeon-Green (mNG) reporter, at the indicated multiplicities of infection (MOI) and the infection efficiency was monitored 48 h later using flow cytometry. Data represent the mean and SD of 3 independent biological replicates. An unpaired *t*-test was applied to assess if the means were significantly different; ****, *p* < 0.0001. **B. Left:** PI4P and GOLPH3 immunofluorescence analysis in A549-ACE2 cells. Cells were non-infected or infected with SARS-CoV-2 at MOI 0.1 and 0.5 and fixed 24 h post-infection prior to PI4P and GOLPH3 staining, as indicated. Images representative from 4 independent experiments are shown. Scale bar is 12 µm. **Right:** The area occupied by GOLPH3 signals was quantified in all cells from one image and divided by the number of nuclei in that image, yielding a mean area per image. Each dot in the graph shows the mean from one image. Colours refer to the 4 independent experiments. The black horizontal line is the mean of all the plotted values. A control incubating cells with PIK93 to inhibit PI4P synthesis and force GOLPH3 signals shrinkage was included. **C. Top:** scheme showing OSBP1’s working mode, which is inhibited by SWG. **Bottom:** A549-ACE2 cells were pre-incubated for 1 h either with DMSO or the indicated concentrations of SWG, infected with SARS-CoV-2(mNG) at the indicated MOIs, and infection efficiency was monitored 48 h later by flow cytometry as in (A). Data represent the mean and SD of 3 independent biological replicates. An unpaired *t*-test was applied to assess if the means were significantly different. **, *p* < 0.01; ***, *p* < 0.001; ****, *p* < 0.0001. **D.** Proximity ligation assays (PLAs) were performed in A549-ACE2 cells infected with SARS-CoV-2 (at the indicated MOIs) or not, using GOLPH3 and PI4P antibodies. Images were acquired using a Zeiss AxioImager Z2 microscope. **Top:** representative images. Scale bar: 12 µm. **Bottom:** the average area covered by puncta per cell was quantified in 3 biological replicates (each attributed a different color). Small dots correspond to individual images while big dots are the mean of each independent experiment, and the horizontal bar is the mean of the means. At least 200 cells were considered per experiment. A one-way paired ANOVA was applied using the three independent means from each experiment to assess the probability that the population means decrease with infection. *, *p* < 0.05; **, *p* < 0.01; ns, non-significant. **E.** GOLPH3 was depleted or not (Control) in A549-ACE2 cells using RNAi (depletion efficiency is shown in Figure S1A), 72 h later cells were infected with SARS-CoV-2(mNG) at the indicated MOIs, and infection efficiency was monitored after 48 h by flow cytometry. Data represent the mean and SD of 3 independent biological replicates. An unpaired *t*-test was applied to assess if the means were significantly different. *, *p* < 0.05. **F.** Similar experiment as in (E) with siRNA-mediated depletion of SAC1 (depletion efficiencies are shown on Figure S1B). *, *p* < 0.05; **, *p* < 0.01; ****, *p* < 0.0001.

To determine whether SARS-CoV-2 requires the PI4P pool specifically associated with the Trans Golgi network, this PI4P pool was monitored using a dedicated immunofluorescence protocol (Venditti *et al*, 2019; Ovejero *et al*, 2023). Upon SARS-CoV-2 infection, the Golgi-resident pool of PI4P, visualized here using a validated anti-PI4P antibody (Kalasova *et al*, 2016), became dispersed throughout the cytoplasm, mostly in the shape of foci (Figure 1B). Furthermore, GOLPH3 signals, used as a marker for the Trans Golgi, shrunk at the highest MOI (Figure 1B), pointing at morphological alterations in this Golgi subdomain.

PI4P is transported from the Trans Golgi to the ER by the protein OSBP1, which, in exchange, extracts one cholesterol moiety from the ER and inserts it at the Trans Golgi (Mesmin *et al*, 2013) (Figure 1C, top). This OSBP1-driven transfer is hijacked by certain +RNA viruses to deviate cholesterol towards the DMVs serving as their replicative niches, using PI4P as a counter-exchange molecule (reviewed in (Roingeard *et al*, 2022)). To assess whether OSBP1-mediated transport was important for SARS-CoV-2 replication, cells were treated with two doses of the OSBP1 specific inhibitor schweinfurthin G (SWG) (Péresse *et al*, 2020) prior to infection, and the infection efficiency was analysed 48 h later. SWG significantly inhibited SARS-CoV-2 replication at the two tested concentrations (Figure 1C, bottom), strongly suggesting a role for OSBP1-mediated transport in SARS-CoV-2 replication.

Next, the proximity between PI4P and GOLPH3, which binds PI4P to maintain a deployed Golgi apparatus morphology (Dippold *et al*, 2009), was analysed 24 h post-infection. GOLPH3 is expected to drop away if PI4P is not available at this location (Dippold *et al*, 2009), thus the potential loss of proximity between these two molecules may represent an indirect readout for PI4P availability. This would be in agreement with the decreased area of GOLPH3-labeled Trans Golgi signals (Figure 1B). Proximity ligation assays (PLA) indeed revealed that signals associated to PI4P-GOLPH3 proximity tended to decrease as the viral input increased (Figure 1D). This suggested that the ability of GOLPH3 to interact with PI4P was impacted upon infection, which could be due to a decrease amount of PI4P or a lack of accessibility. Conversely, siRNA-mediated depletion of GOLPH3, which renders PI4P more accessible, was accompanied by a significantly improved infection efficiency (Figures 1E and S1A). In the same line, depletion of SAC1, the phosphatase that consumes PI4P at the ER once it is translocated there by OSBP1 (Mesmin *et al*, 2013) (Figure 1C, top), also significantly increased SARS-CoV-2 infection efficiency (Figures 1F and S1B). Together, our data show that SARS-CoV-2 benefits from the pool of PI4P emanating from the Trans Golgi and translocated by OSBP1 either to the membranes of the ER and/or to viral DMVs.

### SARS-CoV-2 infection activates the DNA Damage Response kinase ATM in the cytoplasm

We have recently discovered that the pool of PI4P associated to the Trans Golgi serves as a docking platform for one of the master kinases of the DNA Damage Response (DDR), ATM (Ovejero *et al*, 2023). As such, manipulation of cholesterol levels in cellular membranes, of PI4P availability, or of OSBP1 activity, impacts ATM-PI4P interaction (Ovejero *et al*, 2023). Under basal conditions, ATM exists either as a dimer or a multimer (Bakkenist & Kastan, 2003; Wang *et al*, 2016). ATM auto-activation occurs through its auto-phosphorylation, which dismantles the dimer and releases monomers, phosphorylated at serine 1981 (Bakkenist & Kastan, 2003). In line with this, we demonstrated that the use of an ATM inhibitor that stabilizes ATM in its inactive, unphosphorylated form, also stabilizes the PI4P-ATM interaction (Ovejero *et al*, 2023) (scheme in Figure 2A). Considering these previous findings as well as the role of PI4P in SARS-CoV-2 replication, we asked whether SARS-CoV-2 could interfere with the ATM-PI4P interaction. More specifically, given that SARS-CoV-2 hijacks Trans Golgi-associated PI4P, infection could presumably disrupt the ATM-PI4P interaction. Using PLA, we indeed observed a trend towards the loss of ATM-PI4P proximity signals upon infection of A549-ACE2 cells (Figure 2B). Importantly, this decrease seemed antagonized by the ATM inhibitor AZD0156, which stabilizes the ATM-PI4P interaction (Figure 2B).

**Figure 2.**
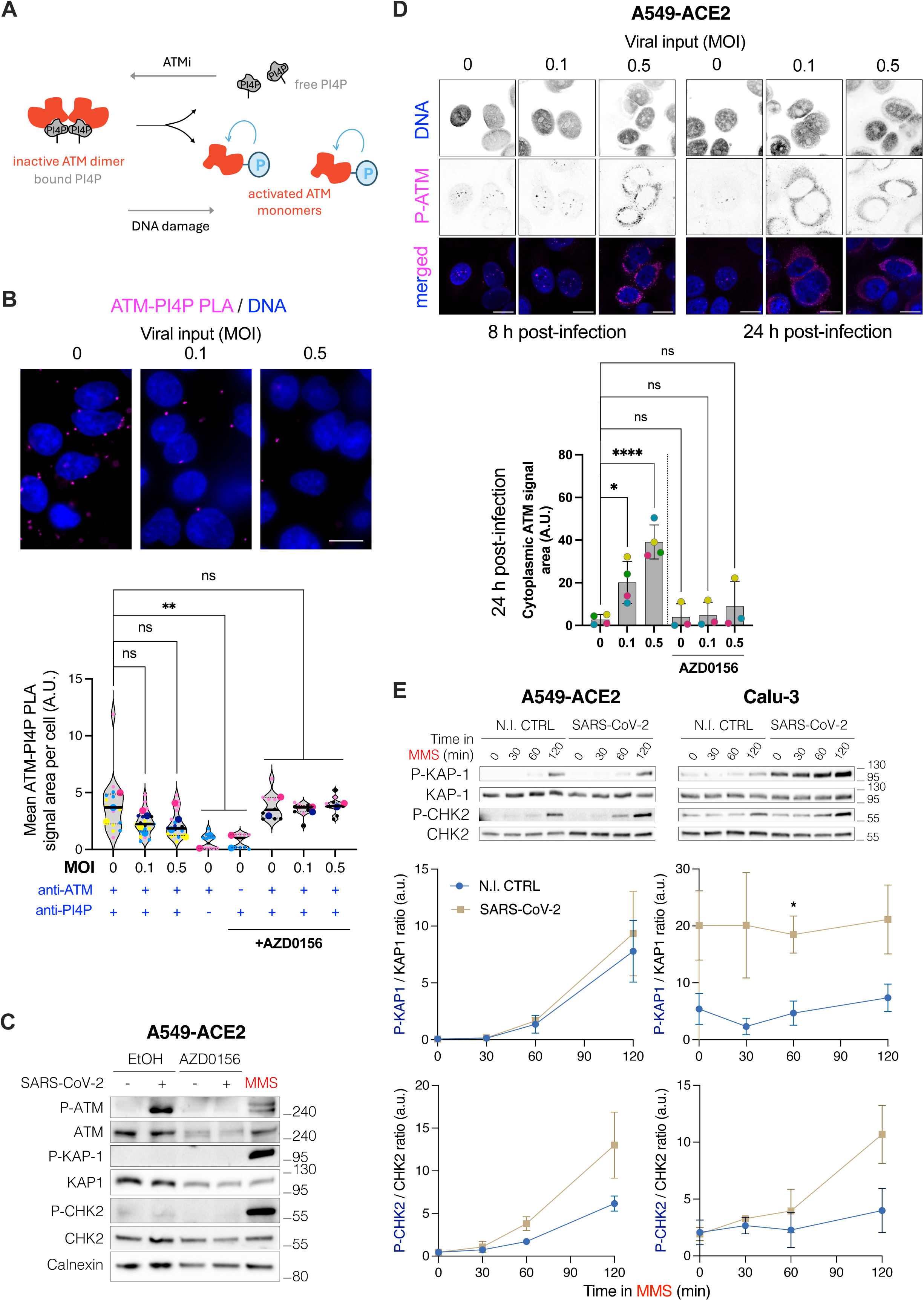
SARS-CoV-2 infection induces ATM activation in the cytoplasm. **A.** Scheme of the master kinase of the DNA Damage Response (DDR) ATM, which is inactive when bound to the pool of PI4P at the Golgi (Ovejero *et al*, 2023). Inactive ATM exists as a dimer, while its activation (by autophosphorylation) leads to its monomerization (Bakkenist & Kastan, 2003) and correlates with its dissociation from PI4P (Ovejero *et al*, 2023). **B.** Proximity ligation assays (PLAs) in A549-ACE2 cells infected or not with SARS-CoV-2 (at the indicated MOIs) and treated or not with AZD0156, using ATM and PI4P antibodies. Images were acquired using a Zeiss AxioImager Z2 microscope. **Top:** representative images, PLA signals are shown in magenta. Scale bar is 12 µm. **Bottom:** the average area covered by puncta per cell was quantified in 3 biological replicates (each attributed a different color). Small dots correspond to individual images while big dots are the mean of each independent experiment, and the horizontal bar is the mean of the means. At least 200 cells were considered per experiment. A one-way unpaired ANOVA was applied using the independent means from each experiment to assess the probability that the population means decrease with infection. ns, non-significant; **, *p* < 0.01. **C.** Western blot analysis of total ATM, autophosphorylated ATM (P-ATM), total KAP-1, phosphorylated KAP-1 (P-KAP-1), total CHK2 and phosphorylated CHK2 (P-CHK2) in A549-ACE2 cells upon infection (at MOI 0.5) and/or AZD0156 treatment, or no treatment. The DNA-damaging agent methyl methanesulfonate (MMS) serves as a positive control for ATM autophosphorylation, and calnexin as a loading control. **D. Top:** Autophosphorylated ATM (P-ATM) was monitored by immunofluorescence analysis in A549-ACE2 cells at two different time points post-infection (8 h and 24 h, as indicated) with two different viral inputs. Representative images acquired with Zeiss LSM880 are shown. Scale bars: 15 µm. **Bottom:** the average area covered by cytoplasmic P-ATM signals was quantified at 24 h post-infection and in the indicated conditions in >200 cells per experiment and in 3 to 4 biological replicates (the mean of each is shown with a different color). The horizontal black line indicates the mean of the means. A one-way unpaired ANOVA was applied using the independent means from each experiment to assess the probability that the population means decrease with infection. ns, non-significant; *, *p* < 0.05; ****, *p* < 0.0001. **E. Top:** Phosphorylation kinetics of ATM targets KAP-1 (chromatin-bound) and CHK2 (soluble, mostly cytoplasmic) assessed by western blots in A549-ACE2 and Calu-3 cells infected with SARS-CoV-2 or not (non-infected control, N.I. CTRL) for 22 h prior to addition of the DNA-damaging agent MMS for up to 2 h, as indicated. Total KAP-1 and CHK2 protein levels are also shown. **Bottom:** Quantifications of the ratio of phosphorylated *versus* total protein signals for both proteins in the indicated cell types and conditions as a function of time since MMS addition. Data show the mean and SEM of three independent experiments. Individual *t*-tests were applied to compare the differences of the means at each timepoint. The single statistically significant difference is indicated, where *, *p* < 0.05.

As this strongly suggested that infection may trigger the release of ATM from its PI4P-bound, resting status, we postulated that this could likely prime its auto-activation by phosphorylation. To assess this, ATM phosphorylation levels were first assessed by western blot. In agreement with the prediction, SARS-CoV-2-infected cells displayed P-ATM signals comparable to those elicited by a classical ATM activator, namely the DNA-damaging agent methyl methanesulfonate (MMS) (Figure 2C). The use of the ATM inhibitor AZD0156 abolished these signals, showing their specificity (Figure 2C). Next, P-ATM localization was assessed by immunofluorescence upon infection. This showed that SARS-CoV-2 elicited a robust accumulation of P-ATM in a time- and viral dose-dependent manner in A549-ACE2 cells (Figure 2D). Of note, in striking contrast with the typical formation of P-ATM foci within the nucleus of cells when these were exposed to MMS (Figure S2A), P-ATM accumulated in the cytoplasm upon infection (Figures 2D and S2A). This phenomenon was observable in all tested model cell lines, whether naturally permissive to SARS-CoV-2 infection (i.e. simian Vero E6 cells, human lung cancer Calu-3 cells) or engineered to express the ACE2 receptor and thereby become permissive (human lung cancer A549-ACE2 and human retinal RPE-1-ACE2 cells) (Figure S2A). Moreover, the P-ATM signals elicited by SARS-CoV-2 were in close apposition, but not co-localizing with, the Trans Golgi marker TGN46 (Figure S2B), in agreement with the release of ATM to the surrounding cytoplasm after PI4P dismantling.

### SARS-CoV-2 infection interferes with ATM-dependent DNA damage signaling

SARS-CoV-2 was recently shown to compromise DNA integrity by provoking the decrease of the protein CHK1, necessary to signal and solve problems during DNA replication (Gioia *et al*, 2023) (of note, this DNA surveillance pathway occurs mostly independently from ATM (Ciccia & Elledge, 2010)). The apparition of the phosphorylated form of the histone H2AX (γH2AX) in infected cells’ nuclei was demonstrated as a downstream event of this deregulation (Gioia *et al*, 2023). Immunofluorescence analyses of infected A549-ACE2 and Vero E6 cells indeed confirmed a modest increase in γH2AX signals upon infection (Figure S2C). However, our data suggested altogether that cytoplasmic ATM activation could be a *bona-fide* consequence of SARS-CoV-2 predation of Golgi-associated PI4P, and therefore unlikely related to downstream consequences of the infection inducing DNA damage. To explore this, the phosphorylation status of two different targets of ATM, strictly chromatin-bound KAP-1 and soluble and mostly cytoplasmic CHK2, were assessed in A549-ACE2 cells. Western blot analysis showed that none of these factors became phosphorylated upon SARS-CoV-2 infection (Figure 2C). This reinforced the view that infection likely elicits ATM phosphorylation by dismantling Trans Golgi-associated PI4P.

To next assess the possibility that pre-activated ATM residing in the cytoplasm could however be reactive to DNA damage arising in parallel to infection, control and infected cells were treated with MMS and the phosphorylation of the two aforementioned ATM targets, KAP-1 and CHK2, was monitored over a 2-hour period (Figure 2E). In the lung cancer cell line Calu-3, KAP-1 was already phosphorylated to some extent at steady state, as typically observed in some cancer cells, and was robustly phosphorylated in infected cells (Figure 2E, right panel), while this was not the case in the other lung cancer cell line A549-ACE2 (as observed before, Figure 2C). In the pre-activated KAP-1 set-up observed in Calu-3 cells, MMS addition did not lead to an increase in phosphorylation. A progressive phosphorylation of KAP-1 could however be monitored in A549-ACE2 cells, which appeared similar in uninfected and infected cells. In contrast, when looking at the activation kinetics of the cytoplasmic target CHK2, a recurrent trend for a faster appearance of P-CHK2 signals was observed in infected A549-ACE2 and Calu-3 cells as compared to non-infected cells (Figure 2E). Thus, SARS-CoV-2 infection seemed to accelerate the phosphorylation kinetics of at least one target of ATM in response to DNA damage. In line with our observation that P-ATM is mostly present in the cytoplasm during SARS-CoV-2 infection, this boost might only apply to cytoplasmic targets.

### Pharmacological inhibition of ATM impairs SARS-CoV-2 replication

As shown in Figure 2B, the ATM inhibitor AZD0156, which prevents ATM auto-activation by phosphorylation and thereby induces a stabilization of ATM interaction with PI4P (Ovejero *et al*, 2023), appeared to stabilize this interaction in the presence of SARS-CoV-2. This raised the possibility that the use of such ATM inhibitors might antagonize SARS-CoV-2 replication. First, immunofluorescence analyses using an antibody specific for viral dsRNA intermediates showed that, both in A549-ACE2 cells (Figure 3A) and in Calu-3 cells (Figure S3A), the ATM inhibitor AZD0156 indeed resulted in a strong decrease in the detection of viral dsRNA replication intermediates. Quantification of viral RNAs (using RdRp RT-qPCR) confirmed a significant inhibitory impact on viral replication, in both cell lines, of AZD0156 (Figure 3B and S3B) and, to a lower extent, of KU55933, a similarly acting ATM inhibitor (Figure S3C). Analysis of the percentage of infected cells using the mNG reporter virus also confirmed the significant antiviral effect of AZD0156 in another model cell line (RPE-1-ACE2 cells) (Figure S3D). This antiviral effect could eventually be overcome by using a high MOI (Figure S3D, MOI 5). Of note, inhibition of other master kinases of the DDR, such as ATR (with VE-821) and DNA-PK (with NU-7026), did not have the same inhibitory effect in two different model cell lines (Figure S3E). This strongly argued that the anti-SARS-CoV-2 effect of ATM inhibitors was presumably related to ATM’s link to Golgi-bound PI4P. In addition, while ATR inhibition had a mild protective effect against infection, in agreement with a previous report (Garcia *et al*, 2021) DNA-PK inhibition was not protective (Figure S3E). Our hypothesis thus stated that ATM inhibition was indirectly antiviral because it promotes tight ATM-PI4P interactions, thus antagonizing viral access to Trans Golgi PI4P pools. An important prediction of this model was that, contrary to effects that could indirectly stem from nuclear DNA damage (where both ATM inhibition and ATM absence would be expected to yield similar phenotypes), ATM removal from the cells should not hamper viral infection efficiency, or might even improve it. In other terms, if the model was correct, ATM inhibition and ATM absence were not expected to have similar consequences. To probe this notion, the impact on SARS-CoV-2 replication of ATM depletion was analysed in in A549-ACE2 cells using either siRNA (Figure S4A) or CRISPR-Cas9 KO (Figure S4B). None of these interventions negatively impacted on SARS-CoV-2 replication (Figure S4C and S4D), reinforcing the model that ATM inhibition could antagonize SARS-CoV-2 infection by restricting viral access to Golgi-bound PI4P.

**Figure 3.**
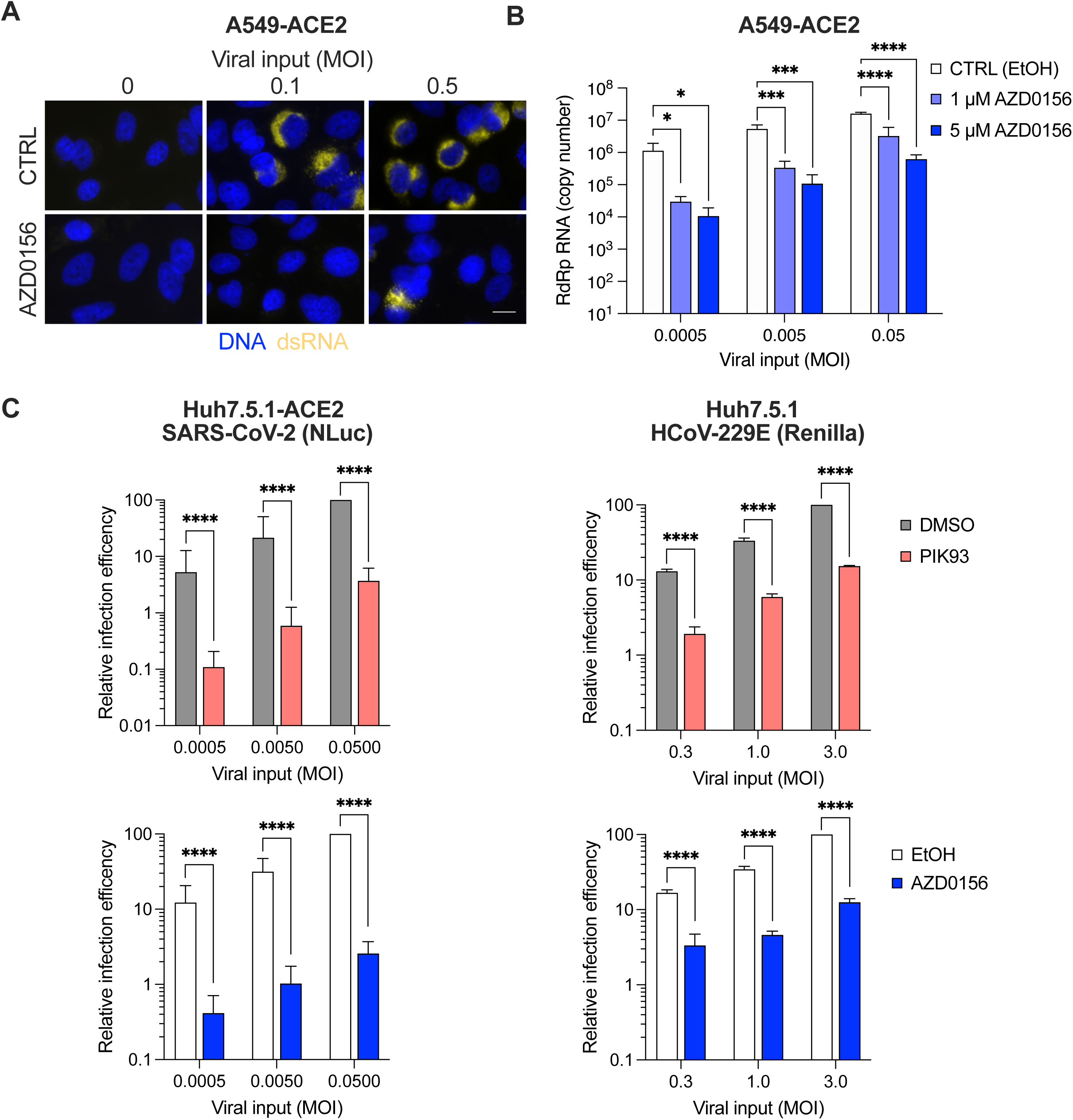
ATM inhibition protects against infection with SARS-CoV-2 and a seasonal coronavirus. **A.** A549-ACE2 cells were incubated either with ethanol as a control or with ATM inhibitor AZD0156 at 5 μM 1 h prior to infection with SARS-CoV-2 at the indicated MOIs for 24 h before cell fixation and dsRNA was detected by immunofluorescence. Images are representative of 3 independent experiments. Images were acquired using a Zeiss AxioImager Z2 microscope. Scale bar: 15 µm. **B.** Viral RNA accumulation upon SARS-CoV-2 infection at the indicated MOIs in the presence or absence (EtOH) of AZD0156 at the indicated concentrations, analysed by RdRp RT-qPCR in A549-ACE2 cells. Mean and SD of 3 independent experiments. One-way ANOVA; *, *p* < 0.05; **, *p* < 0.01; ***, *p* < 0.001; ****, *p* < 0.0001. **C.** Impact of PIK93 (30 μM) or AZD0156 (5 μM) treatments *versus* their respective controls (DMSO or ethanol, EtOH, as indicated) on infection with the indicated MOIs of SARS-CoV-2-Nanoluciferase (NLuc) in Huh7.5.1-ACE2 cells (left) or HCoV-229E-Renilla in Huh7.5.1 cells. Cells were lysed at 24 h post-infection and luciferase activity monitored as a surrogate for replication. Multiple linear regression analysis on log-transformed data; ****, *p* < 0.0001.

Lastly, we wondered whether other coronaviruses could rely on the same pathway and therefore be sensitive to PI4P and ATM inhibition. The impact of pharmacological inhibitors on the aforementioned key actors in PI4P metabolism was tested on the replication efficiency of a seasonal alphacoronavirus, HCoV-229E, in parallel to SARS-CoV-2, using luciferase-expressing viruses in the Huh7.5.1 model cell line (ectopically expressing ACE2 or not) (Figure 3C). Inhibition of PI4P synthesis with PIK93 and of ATM with AZD0156 had a similar impact on HCoV-229E than on SARS-CoV-2, suggesting that PI4P predation could be a conserved feature across human coronaviruses.

## Discussion

In this study, we have revealed that SARS-CoV-2 diverts PI4P available at the Trans Golgi network by exploiting the OSBP1 transporter. This likely serves to develop its replicating niche, the DMVs, either by direct transfer, or *via* passage of PI4P molecules through the ER. Consequently, PI4P is less available at the Golgi for its natural binders, which include GOLPH3, a factor implicated in Golgi morphology, and ATM, a master upstream kinase of the DDR. This discovery has functional consequences: we characterized in detail that the loss of the ATM-PI4P interaction fosters the auto-phosphorylation activity of ATM, which we discovered becomes massively enriched in the host cell’s cytoplasm. This unveiled the reciprocal notion that inhibiting ATM’s auto-phosphorylation can stabilize the ATM-PI4P couple, consequently antagonizing SARS-CoV-2 infection. These concepts hold true in multiple model cell lines and apply to at least another human coronavirus. Moreover, we found that ATM activation as a consequence of infection primes an acceleration of its signalling activities in response to DNA damage.

### New light on SARS-CoV-2 infection

We find that SARS-CoV-2 infection leads to PI4P consumption by co-opting the OSBP1 transporter. For this to work, OSBP1 necessitates continuous fuelling with ER-based cholesterol moieties, which are used to pay against transported PI4P molecules (Mesmin *et al*, 2013). We reveal its similar dependence on continuous PI4P synthesis (Figure 1A) and on OSBP1 itself (Figure 1C). The latter is in agreement with a recent report showing an impact of OSBP inhibition on coronavirus replication (Ma-Lauer *et al*, 2024). PI4P could serve to develop the virus’s replicating niche, and be transported either by following a canonical route from the Golgi to the ER then be deviated to the viral DMVs; alternatively, OSBP1 could transfer PI4P directly from the Golgi to the DMVs. In both cases, DMVs would be expected to provide cholesterol in exchange. This need would be in agreement with the numerous reports that highlight SARS-CoV-2’s dependency on cholesterol. There is precedent for the PI4P-cholesterol counterflow mechanism being hijacked for the benefit of viral replication. For instance, other RNA viruses, the picornavirus poliovirus (Arita, 2014), the flavivirus HCV (Wang *et al*, 2014), or rhinoviruses (Roulin *et al*, 2014), all hijack OSBP to structure their DMVs.

Concerning SARS-CoV-2 specifically, nsp4 was proposed to interact both with OSBP and with the PI4-kinase PI4KIIIβ (St-Germain *et al*, 2020). This points at this viral factor as potentially responsible for the co-opting mechanism we describe here. This provocatively meets the fact that the simple expression of nsp3, nsp4, and nsp6 viral proteins, all of which possess transmembrane and cytosolic domains, suffices to engage ER transitions yielding DMVs precursors (Angelini *et al*, 2013). This had led us to propose that nsp3-4-6 viral proteins instruct the dimerization of ER membranes that would initiate a lamellar-to-cubic bilayer transition able to support DMVs structuration (Moriel-Carretero, 2020).

Functionally, the PI4P-rich lipid microenvironment within the DMVs is key to promote RNA replication of enteroviruses and the flavivirus HCV, as PI4P binds to RNA-dependent RNA polymerases and regulate their activity (Hsu *et al*, 2010). Similarly, PI4P-rich DMVs are generated that facilitate replication of the viral genome in the context of after picornavirus Seneca Virus A infection (Zhao *et al*, 2023a). In the case of SARS-CoV-2, N has been shown to interact with PI4P and other phosphoinositides (Dutta *et al*, 2024).

### The impact of SARS-CoV-2 infection on Golgi function

Our work implies a direct impact of SARS-CoV-2 infection on the Trans Golgi structure and/or function because of PI4P deprivation. First, PI4P exhaustion or monopolization by SARS-CoV-2 limits access to its natural binders. Such is the case of GOLPH3, as our PLA experiment shows that the interaction between PI4P and GOLPH3 decreases with infection (Figure 1D). By connecting Golgi-bound PI4P with the cytoskeleton, GOLPH3 maintains Golgi morphology (Dippold *et al*, 2009). Moreover, the accurate PI4P levels granted by intact OSBP1 activity promote the formation of lipid gradients along the Trans Golgi that shape cargo sorting. This is essential in polarized epithelial cells for the dispatch and trafficking of numerous plasma membrane cargo proteins with apical or basolateral localization (Kovács *et al*, 2024). Last, the fine control of specific PI4P profiles in the Trans Golgi dictates the profiles of glycosylation of resident Golgi proteins ((Bankaitis *et al*, 2012); pre-print (Sasaki *et al*, 2023)), and its alteration provokes the activation of the Golgi Stress Response (reviewed in (Sasaki & Yoshida, 2019)).

Despite these notions suggesting that SARS-CoV-2 infection would lead to the dismantling of Golgi functions, it must be remembered that intra-Golgi activities are key to support the final stages of replication aiming at viral egress. Hence, coronaviral Envelope (E) proteins must undergo a Golgi-to-lysosome trafficking, where E acts to deacidify lysosomes prior to viral release (Pearson *et al*, 2024). It is therefore more likely that PI4P predation ends up rewiring Golgi functions to further serve late viral cycle steps, rather than leading to the disintegration of the Golgi.

### SARS-CoV-2 and the DDR

SARS-CoV-2 infection has been reported to challenge genome integrity by interfering with processes susceptible of activating the ATR branch of the DDR. First, in simian Vero E6 cells, the ATR axis was activated following the infection-induced downregulation of the telomere-protecting factor TRF2, as this results in unprotected telomeres (Victor *et al*, 2021). Second, in infected model cell lines and mice tissues, as well as in nasal and lung samples from COVID-19 patients, the downstream effector of the ATR signalling pathway CHK1 was subjected to degradation. This prevented the host cells that underwent DNA replication to ensure replication fork protection and repair in the case of encountering difficulties, a phenomenon known as replicative stress (Gioia *et al*, 2023). Another coronavirus, Infectious Bronchitis Virus (IBV), elicited DNA replicative stress through interaction with polymerase delta, impairing the correct progression of cells through the S phase (Xu *et al*, 2011). This study openly proposed that activation of the host’s DDR represents a mechanism exploited by coronaviruses to induce cell cycle arrest, thus redirecting resources to viral replication instead of that of the host. In support of this interpretation, suppression of the ATR kinase activity either by chemical inhibition or *via* siRNA-mediated knockdown inhibited IBV replication. More recently, pharmacological inhibition of ATR with berzosertib was also shown to decrease SARS-CoV-2, SARS-CoV-1 and MERS-CoV replication (Garcia *et al*, 2021). In line with this, we observed a modest inhibition of SARS-CoV-2 with ATR inhibitor VE-821.

A very different picture emerges for the other master kinase of the DDR, ATM. To our knowledge, there were no previous reports linking ATM and SARS-CoV-2 infection, apart from a study that observed autophosphorylation of ATM upon infection (Gioia *et al*, 2023), and another one reporting an interaction between ORF7a and ATM (Stukalov *et al*, 2021). The first work assumed this was due to infection-triggered CHK1 degradation and host cell’s replicative stress, and was however not accompanied by the phosphorylation of ATM’s main downstream target CHK2. This observation was not pursued further. We detected P-ATM upon SARS-CoV-2 infection by western blot, and confirmed this was not accompanied by a matching detection of P-CHK2. The additional use of immunofluorescence analysis allowed us to uncover an atypical profile, with a strong P-ATM signal in the cytoplasm. Together, our data strongly pointed towards a DNA damage-independent cause as the one responsible for ATM activation. Indeed, we discovered that this activation is a collateral consequence of SARS-CoV-2 predating PI4P, to which ATM is bound at the Trans Golgi during its resting state (Ovejero *et al*, 2023) (Figures 2 & S2). Reinforcing this notion, the stabilization of the PI4P-ATM association by ATM inhibitors greatly prevented viral replication (Figures 3 & S3). Importantly, if ATM activation stemmed from DNA damage signals, both the chemical inhibition of ATM and removing the protein would hamper viral replication, as is the case for ATR (see above). Yet, ATM depletion did not affect SARS-CoV-2 (Figure S4). Thus, we reveal here an unexpected and DNA damage-independent means by which a master kinase of the DDR becomes activated upon coronavirus infection.

As mentioned, the activation of ATM in the cytoplasm did not concur with a parallel activation of its targets, whether CHK2 or KAP-1. This agrees with the knowledge that, beyond its own auto-phosphorylation, an appropriate DNA signal (i.e., DNA extremities looking like a double strand break) needs also to be present to stimulate ATM (Ismail *et al*, 2005). In line with this, only when actual DNA damage was provoked using a genotoxin could we detect the phosphorylation of such ATM targets (Figure 2E). Importantly, though, the cytoplasmic existence of an infection-induced, pre-activated pool of ATM led to an accelerated kinetics in the phosphorylation of (cytoplasmic) CHK2 effector (Figure 2E). In contrast, the kinetics of activation of the chromatin-bound effector KAP-1 remained unchanged, likely because ATM needs for this to access the nucleus, as in standard DNA damage scenarios (Figure 2E). This observation might have important implications on our understanding of the known strong susceptibility of cancer patients undergoing chemotherapy. Indeed, it has been reported that cancer, and chemotherapies, are susceptibility factors for developing severe or even lethal COVID-19 (Williamson *et al*, 2020); (Albiges *et al*, 2020). Activated CHK2 is responsible for maintaining the cells arrested in a pro-apoptotic status, and also contributes to the secretion of pro-inflammatory interleukins 6 and 8 (Rodier *et al*, 2009), both part of the cytokine storm observed in severe cases of COVID-19 (Hu *et al*, 2021; Elbadawy *et al*, 2023). It is therefore likely that, upon chemo- or radiotherapeutic challenge, cancer patients endure an enforced ATM-CHK2 axis activation that aggravates the already toxic nature of this treatment. Hence, we propose that the cytokine storm that worsens patients’ prognosis might, at least in part, derive from the molecular mechanism we unveil in this work.

In this set-up, the potential use of ATM inhibitors not only as antivirals, but also as anti-inflammatory agents, might be worth considering. It has been proposed, as noted above, that inhibition of the DDR kinase ATR may hold therapeutic benefit against SARS-CoV-2 replication (Garcia *et al*, 2021). However, as we show in two different cell types (Figure S3), at an identical concentration, ATM inhibition emerges as clearly more powerful.

In sum, we have uncovered an unconventional mode of activation of the DDR kinase ATM that reveals both a fundamental aspect of SARS-CoV-2 infection, namely the exploitation of Golgi-associated PI4P, and an unforeseen strategy to help tackle COVID-19 by inhibiting one of the main axes of the DDR.

## Materials and methods

### Plasmids

The plasmids used to produce HIV-1 based lentiviral vector (p8.91 coding HIV-1 Gag-Pol, pMD.G coding VSV Glycoprotein, G) with ACE2 coding minigenome (pRRL.sin.cPPT.SFFV/ACE2.WPRE; Addgene 145842) have been described (Rebendenne *et al*, 2021; Naldini *et al*, 1996). The pLX_311-Cas9 and LentiGuide-Puro vectors were gifts from John Doench and Feng Zhang, respectively (Doench *et al*, 2014; Sanjana *et al*, 2014) (Addgene 96924 and 52963) and we have described before the LentiGuide-Puro-CTRLg1 and g2 (Rebendenne *et al*, 2021). Guide RNA coding oligonucleotides were annealed and ligated into BsmBI-digested LentiGuide-Puro vector, as described (Addgene). The primer sequences used were as follow: gATM1-Fwd 5’-caccgTCTACCCCAACAGCGACATG, gATM1-Rev 5’-aaacCATGTCGCTGTTGGGGTAGAc, gATM2-Fwd 5’-caccgGACCTACCTGAATAACACAC, gATM2-Rev 5’-aaacGTGTGTTATTCAGGTAGGTCc.

### Cell culture

Human HEK293T, A549, RPE-1, and Calu-3, and simian Vero E6 cells were maintained in complete Dulbecco’s modified Eagle medium (DMEM) (Gibco) supplemented with 10% foetal bovine serum and penicillin/streptomycin. Calu-3 cells were obtained from American Type Culture Collection (ATCC); HEK293T, A549, Vero E6 cells were gifts from Michael Malim’s lab, Wendy Barclay’s lab, and from the CEMIPAI facility, respectively. RPE-1 cells were a kind gift from Urszula Hibner and were authenticated by ATCC STR profiling. All cell lines were regularly screened for the absence of mycoplasma contamination. A549 and RPE-1 cells stably expressing ACE2 were generated by transduction with RRL.sin.cPPT.SFFV.WPRE containing-lentiviral vector.

For CRISPR-Cas9-mediated gene disruption, cells stably expressing Cas9 were first generated by transduction with LX_311-Cas9 followed by blasticidin selection at 10 µg/ml. Cas9 activity was checked using the XPR_047 assay (a gift from David Root, Addgene 107145) and was >90%. The cells were then transduced with guide RNA expressing LentiGuide-Puro and selected with antibiotics for at least 10 days.

When indicated, PI4P synthesis inhibitor PIK93 (10 or 30 µM) ATM inhibors AZD0156 (1 or 5 µM) or ATR inhibitor VE-821 (5 µM), DNA-PKcs inhibitors NU-7026 (0.5 ou 5 µM), remdesivir (5 µM), SWG (xM) or similar volumes of their diluents (DMSO or ethanol) were added 1 h prior to infection, and the molecules left over the course of infection. When used as a control, MMS was added at 0.005% for the indicated duration.

### Lentiviral production and trasnduction

Lentiviral vector stocks were obtained by polyethylenimine (PEI; for LentiGuides) or Lipofectamine 3000 (Thermo Scientific; for ACE2)-mediated multiple transfections of 293T cells in 6-well plates with vectors expressing Gag-Pol, the viral minigenome, and VSV G at a ratio of 1:1:0.5. The culture medium was changed 6 h post-transfection, and vector containing supernatants harvested 36 h later, filtered and used directly or stored at −80°C.

### RNAi

siRNA transfections were performed with X-tremeGENE™ 360 (Merck) according to the manufacturer’s instructions using 40 nM final of the different siRNAs.

### SARS-CoV-2 production and infection

The BetaCoV/France/IDF0372/2020 isolate was supplied by Pr. Sylvie van der Werf and the National Reference Centre for Respiratory Viruses hosted by Institut Pasteur (Paris, France). The patient sample from which strain BetaCoV/France/IDF0372/2020 was isolated was provided by Dr. X. Lescure and Pr. Y. Yazdanpanah from the Bichat Hospital, Paris, France. The mNeonGreen (mNG) (Xie *et al*, 2020a) and Nanoluciferase (NLuc) (Xie *et al*, 2020b) reporter SARS-COV-2 were based on 2019-nCoV/USA_WA1/2020 isolated from the first reported SARS-CoV-2 case in the USA, and provided through World Reference Center for Emerging Viruses and Arboviruses (WRCEVA), and UTMB investigator, Dr. Pei Yong Shi (Xie *et al*, 2020b, 2020a). WT, mNG and NLuc reporter SARS-CoV-2 were amplified in Vero E6 cells (MOI 0.005) in serum-free media. The supernatant was harvested at 48 h-72 h post infection when cytopathic effects were observed, cell debris were removed by centrifugation, and aliquots frozen down at −80°C. Viral supernatants were titrated by plaque assays in Vero E6 cells. Typical titers were 3.10^6^-3.10^7^ plaque forming units (PFU)/ml. HCoV-229E-Renilla was a gift from Volker Thiel (van den Worm *et al*, 2012) and was amplified for 5-7 days at 33°C in Huh7.5.1 cells in 5% FCS-containing DMEM. Viral stocks were harvested when cells showed >50% CPEs. Viruses were titrated through TCID_50_ in Huh7.5.1 cells and typical titers were 1-2.10^9^ TCID_50_/mL. Simian Vero E6 and human cell infections with coronaviruses were performed at the indicated multiplicity of infection (MOIs; as calculated from titers in Vero E6 or Huh7.5.1 cells) in 5% serum-containing DMEM. At the indicated time post-infections, the cells were trypsinized, fixed in PBS1X-2% PFA and the percentage of cells expressing mNG was scored by flow cytometry using a NovoCyte^TM^ (ACEA Biosciences Inc.), or the cells were lysed in Lysis buffer and luciferase activity measured (Promega), or lysed in RLT buffer (Qiagen) followed by RNA extraction and RT-qPCR analysis.

### Quantification of mRNA expression

3-5 × 10^5^ cells were harvested and total RNA was extracted using the RNeasy kit (Qiagen) employing on-column DNase treatment, according to the manufacturer’s instructions. 125 ng cellular RNAs were used to generate cDNAs. The cDNAs were analysed by qPCR using published RdRp primers and probe (Corman et al., 2020), as follow: RdRp_for 5’-GTGARATGGTCATGTGTGGCGG-3’, RdRp_rev 5’-CAAATGTTAAAAACACTATTAGCATA-3’ RdRp_probe 5’-FAM-CAGGTGGAACCTCATCAGGAGATGC-TAMRA-3’). qPCR reactions were performed in triplicate, in universal PCR master mix using 900 nM of each primer and 250 nM probe or the indicated Taqmans. After 10 min at 95°C, reactions were cycled through 15 s at 95°C followed by 1 min at 60°C for 40 repeats. Triplicate reactions were run according to the manufacturer’s instructions using a ViiA7 Real Time PCR system (ThermoFisher Scientific). pRdRp (which contains an RdRp fragment amplified from SARS-CoV-2 infected cell RNAs using primers RdRp_for and RdRp_rev and cloned into pPCR-Blunt II-TOPO, (Rebendenne *et al*, 2021)) was diluted in 20 ng/ml salmon sperm DNA to generate a standard curve to calculate relative cDNA copy numbers and confirm the assay linearity (detection limit: 10 molecules of RdRp per reaction).

### Reagents

Methyl methanesulfonate (129925, Sigma-Aldrich),
DAPI (D9542, Sigma-Aldrich),
cOmplete protease inhibitor cocktail (11836170001, Roche),
Halt Phosphatase inhibitor cocktail (78420 ThermoFisher) and
ProLong (P36930, ThermoFisher),
AZD0156 (HY-100016-5mg, Clinisciences),
KU-55933 (M1801-10mg, Clinisciences),
VE-821 (HY-14731, Clinisciences),
PIK93 (HY-12046, Clinisciences),
NU-7026 (A12752-10; Clinisciences),
Remdesivir (T7766-5mg, TargetMol, US)
schweinfurthin G (SWG) was a kind gift from Bruno Mesmin and Bruno Antonny.
siControl was the universal negative control siRNA #1 (SIC001, Sigma-Aldrich), and the remaining siRNAs were also from Sigma-Aldrich:
siATM#1 (SASI_Hs01_00093615, 5’-CUUAGCAGGAGGUGUAAAU[dT][dT]-3’);
siATM#2 (SASI_Hs01_00093616, 5’-CCCAUUACUAGACUACGAA[dT][dT]-3’);
siATM#3 (SASI_Hs01_00093617, 5’-GUUACAACCCAUUACUAGA[dT][dT]-3’);
siGOLPH3#1 (SASI_Hs02_00355527, 5’-GUCUGAAGGCCAACACCAA[dT][dT]-3’);
siSAC1#1 (SASI_Hs01_00199997, 5’-CUCAUUUGGGACUUAUAAU[dT][dT]-3’);
siSAC1#2 (SASI_Hs01_00199998, 5’-GGUUUGUAUGGAAUGGUCA[dT][dT]-3’);
siSAC1#3 (SASI_Hs01_00199999, 5’-CAAUUGCAUGGAUUGUCUA[dT][dT]-3’);

### Antibodies

<u>Primary antibodies:</u>

anti-calnexin (610523, BD transduction Laboratories, WB at 1/ 1,000),
anti-P-Thr68-CHK2 (2661S, Cell Signaling, WB at 1/1,000),
antitotal CHK2 (05-649, Millipore, WB at 1/1,000),
anti-P-Ser1981-ATM (4526, Cell Signaling, IF at 1/200, WB at 1/1,000)
antitotal ATM (GTX132147, GenTex, IF at 1/200, PLA at 1/2,000),
anti-KAP-1 (BETA300-274A, Bethyl, WB at 1/2,000),
antitotal KAP-1 (BETA300-274A, Bethyl, WB at 1/2,000)
anti-PI4P (Z-P004; Echelon Biosciences, PLA at 1/ 2,000; IF at 1/1,000),
anti-P-Ser824-KAP-1 (BETA300-767A, Bethyl, WB 1/2,000),
anti-P-Ser139-H2AX (05–636, Sigma-Aldrich, IF at 1/500),
anti-GOLPH3 (SAB4200341, Sigma-Aldrich, IF at 1/300, WB at 1/ 2,000; PLA at 1/ 2,000),
anti-TGN46 (AHP500G, Bio-Rad, IF at 1/100).
anti-SACM1L (anti-SAC1) (13033-1-AP, ProteinTech Group)
anti-dsRNA J2 (10010200, Scicons, IF 1/1000)
anti-SARS-CoV-2 N (A2060-50, Biovision, IF 1/500)

<u>Secondary antibodies:</u>

<u>for WB:</u>

goat anti-rabbit (A0545, Sigma-Aldrich) at 1/5,000,
rabbit anti-mouse (A9044, Sigma-Aldrich) at 1/ 5,000.

<u>for IF:</u>

DyLight 488 donkey anti-rabbit (406404, Biolegend) at 1/500,
DyLight 649 goat anti-mouse (405312, Biolegend) at 1/500,
anti-sheep Alexa 546 (discontinued, formerly A21088; ThermoFisher) at 1/500.

### Western blot

All the samples were lysed with high-salt buffer (50 mM Tris pH 7.5, 300 mM NaCl, 1% Triton-x-100, protease inhibitors (11836170001, Roche), and Pierce Phosphatase inhibitor cocktail (A32957, ThermoScientific); 90µL for one well of a 24 well-plate) for 10 min on ice with frequent vortexing. Samples were cleared for 10 min at 17,000 g at 4°C, and supernatants quantified using the Pierce^TM^ BCA kit (10741395, ThermoFisher). For western blot, 20-30 µg of whole-cell extracts were loaded in SDS-PAGE and migrated 1h30 at 150V (migration buffer: 25 mM Tris, 200 mM Glycine, 0.1% SDS, for P-ATM detection: 50 mM Tris, 50 mM Tricine, 0.1% SDS). The proteins were transferred to a nitrocellulose membrane with semi-dry system (TransBlot BioRad), or liquid transfer over-night at 4°C at 30 V (transfer buffer: 25 mM Tris, 200 mM Glycine, 20% methanol). Transfer is followed by immunoblotting using primary antibodies then with HRP-coupled secondary antibodies. Detection is done using Super Signal West Pico PLUS ECL (15626144, Thermo Scientific) and acquisition with Imager P2 (Amersham).

### Immunofluorescence

Cells were fixed either using 4% PFA for 20 min or 100% cold methanol for 10min in the case of downstream P-ATM detection. Cells fixed on coverslips were washed once with 1× PBS and permeabilized with 0.2% Triton/1× PBS for 10 min, or with 100 µM Digitonin/1× PBS for 5 min in the case of downstream PI(4)P detection. Then saturated with 3% BSA/1× PBS for 30 min. Coverslips were incubated with primary antibodies diluted in 3% BSA/1× PBS for 1h and then washed 3 times with 1× PBS under gentle shaking. Coverslips were further incubated with secondary antibodies diluted in 3% BSA/1× PBS for 1h while protecting them from light from this point on, then washed 3 times with 1× PBS under gentle shaking, then incubated for 10 min with DAPI (1 µg/ml) diluted in H_2_O, and washed three times with H_2_O. Finally, coverslips were allowed to dry and mounted using ProLong Gold Antifade and then left to dry overnight at room temperature in the darkness.

### Proximity ligation assays

Cells were fixed with 4% PFA/PBS during 20 min at room temperature. Subsequently, every step took place in a humid chamber using the PLA reagents from Sigma-Aldrich. Cells were permeabilized with 0.5% Triton/PBS for 10 min and incubated with Blocking Solution from the kit for 1 h. Primary antibodies (anti-PI4P and anti-ATM or anti-GOLPH3) were diluted at 1:2,000 in the Antibodies Diluent from the kit and incubated overnight at 4°C onto the coverslips. For technical controls, one of these primary antibodies was omitted at a time. PLA minus and plus probes were next mixed and incubated in the Blocking Solution for 20 min as specified by the supplier, then added onto the coverslips, and incubated for 1 h at 37°C. Coverslips were washed twice with Buffer A (150 mM NaCl; 10 mM Tris; 0.05% Tween-20; pH 7.4), then incubated with Ligation Mix (1:40 ligase; 1:5 ligase buffer) 30 min at 37°C, washed again twice with Buffer A, and then incubated with Polymerization Mix (1:80 polymerase; 1:5 polymerase buffer) 1 h at 37°C. Last, coverslips were washed with Buffer B (100 mM NaCl; 200 mM Tris; pH 7.5) and then with 0.01× Buffer B. The coverslips were finally mounted with Duolink in situ Mounting Medium with DAPI and then sealed with nail polish.

### Microscopy, image quantification and plots

Acquisition of immunofluorescence or PLA signal were done using epifluorescence microscopy with an upright microscope (Zeiss AxioImager Z2), at x40 magnification. Confocal images were acquired using an LSM880 microscope (Zeiss) coupled to an AIRYSCAN module, with a 63x lens. AIRYSCAN pre-processing was performed using the ZEN Black software (ZEISS) and post-processing was performed using FIJI. Signals were detected using a FIJI macro subtracting the background, setting the threshold to create a mask, applying the Watershed algorithm, and then analysing particles. The total signal area from one image was divided by the number of nuclei in that image, thus yielding the mean signal area per image. All the individual values obtained from different images are plotted as a dot. All dots belonging to a similar, independent experiment share the same colour. For each experiment, a big dot indicates the mean the smaller dots, that is, the mean per experiment. Horizontal bars indicate the mean of the independent experiments.

### Statistical analysis

GraphPad Prism was used to plot all the graphs and to statistically analyse the data. For statistical analyses, the most relevant test was chosen depending on the nature of the data and these are clearly stated in the corresponding figure legends. When nothing is indicated on the graphs, differences are not significant (ns).

### Biosafety

Experiments with SARS-CoV-2 were performed in a BSL-3 laboratory, and experiments with HCoV-229E in BSL-2 laboratories, following safety and security protocols approved by the risk prevention services of CNRS.

## Abbreviations

ACE2: Angiotensin-Converting Enzyme 2
ATM: Ataxia-Telangiectasia-Mutated
ATR: Ataxia-Telangiectasia and Rad3-related
COVID-19: COronaVirus-Induced Disease 19
DDR: DNA Damage Response
DMV: Double Membrane Vesicle
ER: Endoplasmic Reticulum
HCV: Hepatitis C Virus
MMS: Methyl MethaneSulfonate
MOI: Multiplicity Of Infection
N: Nucleoprotein
OSBP1: OxySterol-Binding Protein 1
PI4P: Phosphatidyl-inositol-4-Phosphate
PLA: Proximity Ligation Assay
SARS-CoV-2: Severe Acute Respiratory Syndrome Coronavirus 2
SWG: schweinfurthin G
RT-qPCR: Reverse Transcription - quantitative Polymerase Chain Reaction

## Disclosure and competing interests statement

The authors declare that have no conflict of interest.

## Authors’ contributions

In addition to the CRediT author contributions, the contributions in detail are: M.M. and C.G. designed the study, analysed the data and wrote the manuscript. A.R., A.L.C.V., O.M. and C.G. carried out the SARS-CoV-2 experiments, C.S. performed the immunoblotting, PLAs, immunofluorescences and all microscopy analyses except for the Airyscan analysis, performed by J.M. B.B. performed the HCoV-229E (and SARS-CoV-2 NLuc) experiments. All authors have read and approved the manuscript.

## Acknowledgements

We are grateful to colleagues sharing reagents (see section “Material & Methods”) and to our funding sources. This work was supported by a Flash Grant Cancer/COVID from Fondation ARC, France (to M.M.), the Fondation CNRS (to C.G.), the European Research Council (ERC) under the European Union’s Horizon 2020 research and innovation programme (grant agreement 759226, ANTIViR, to C.G.), a 3-year PhD studentship from the Ministry of Higher Education and Research (to A.R.), a fourth year PhD funding from the Fondation pour la Recherche Médicale (FRM grant number [FDT202106013175], to J. M.), institutional funds from Centre National de la Recherche Scientifique (CNRS) and Montpellier University. Finally, the authors acknowledge the BSL-3 facility CEMIPAI (UAR 3725 CNRS Montpellier University) and the imaging facility MRI, a member of the national infrastructure France-BioImaging supported by the French National Research Agency (ANR-10-INBS-04).

---

**Figure S1.**
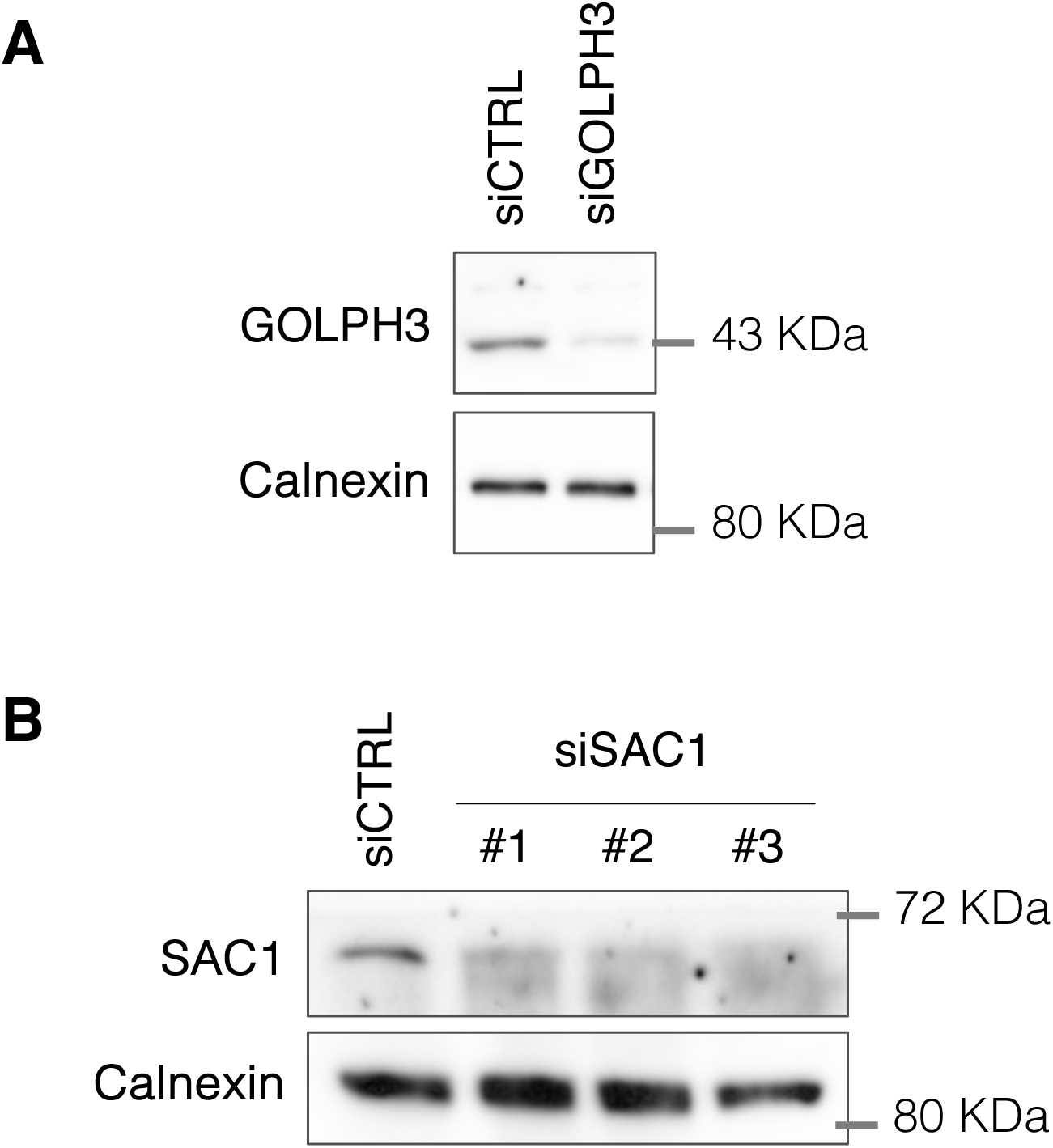
Controls of protein depletions for experiments shown in Figure 1. **A** and **B.** Western blots showing GOLPH3 (**A**) or SAC1 (**B**) depletion 72 h post-transfection with siRNA targeting GOLPH3 or SAC1, as indicated, in comparison to control siRNA (siCTRL). Calnexin serves as a loading control. Molecular sizes are indicated in KDa.

**Figure S2.**
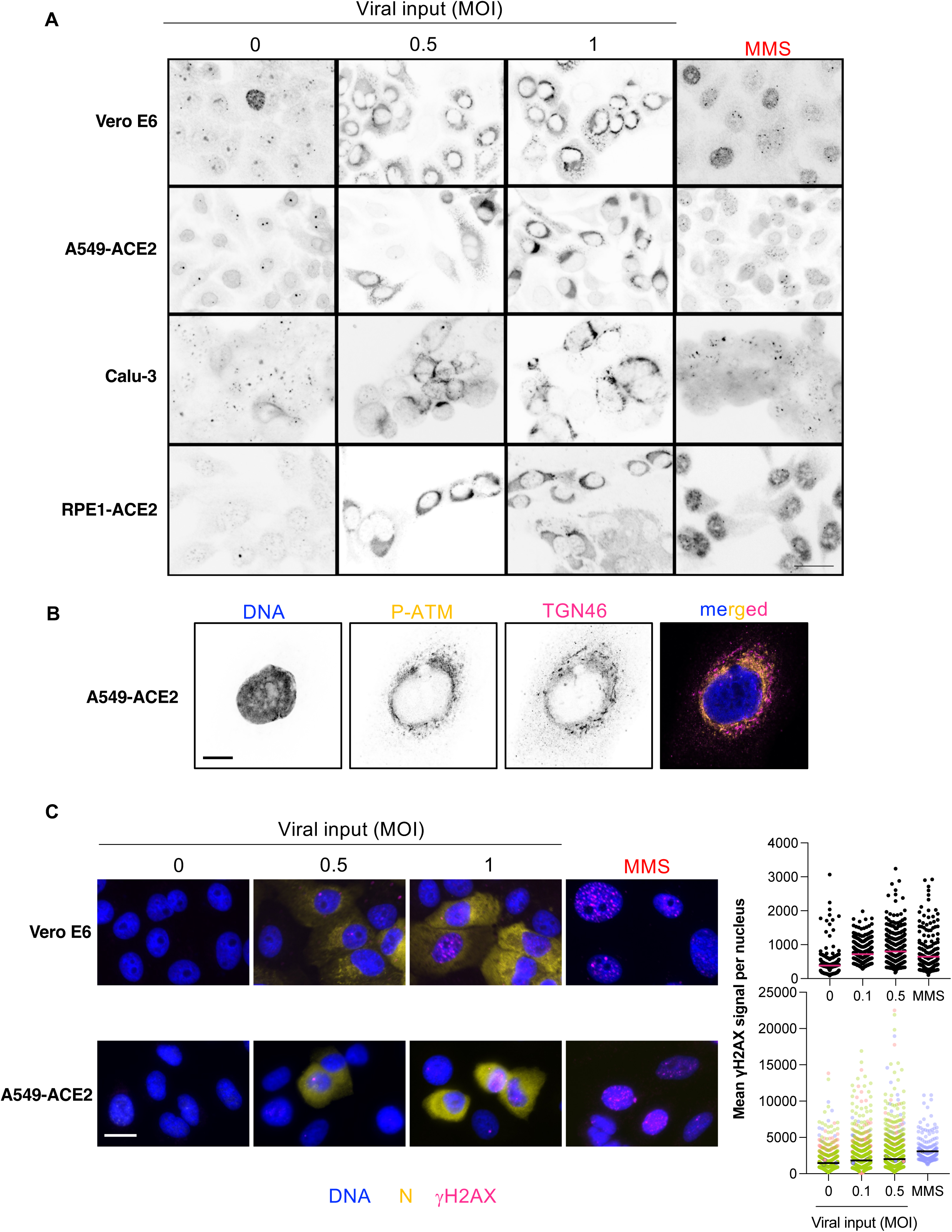
ATM activation in the cytoplasm upon SARS-CoV-2 infection is observed across cell types. **A.** Autophosphorylated ATM (P-ATM) was monitored by immunofluorescence at 24 h post-infection with the two indicated viral doses of SARS-CoV-2, in the 4 indicated cell lines. Scale bar: 30 μm. MMS was included as a treatment generating P-ATM signals in the nucleus. **B.** Immunofluorescence of A549-ACE2 cells 24 h post-infection with SARS-CoV-2 showing that P-ATM signals are in proximity to, but do not overlap with, Trans Golgi signals, as assessed using the marker TGN46. Scale bar: 10 μm. **C. Left:** Representative images of Vero E6 (top) and A549-ACE2 (bottom) cells immunostained 24 h after infection with SARS-CoV-2 (or not), to assess the abundance of the phosphorylated form of H2AX ([H2AX, magenta) in infected (N^+^ cells, yellow) *versus* non-infected cells; nuclei are shown in blue. MMS serves as a positive control. Scale bar: 15 μm. **Right:** the graphs show individual dots representing the mean [H2AX signal per nucleus expressed in arbitrary units. For Vero E6, only one experiment was performed (black color). For A549-ACE2 cells, three experiments were performed, each indicated by a different color. The horizontal lines represent the mean value of all the dots in each cloud.

**Figure S3.**
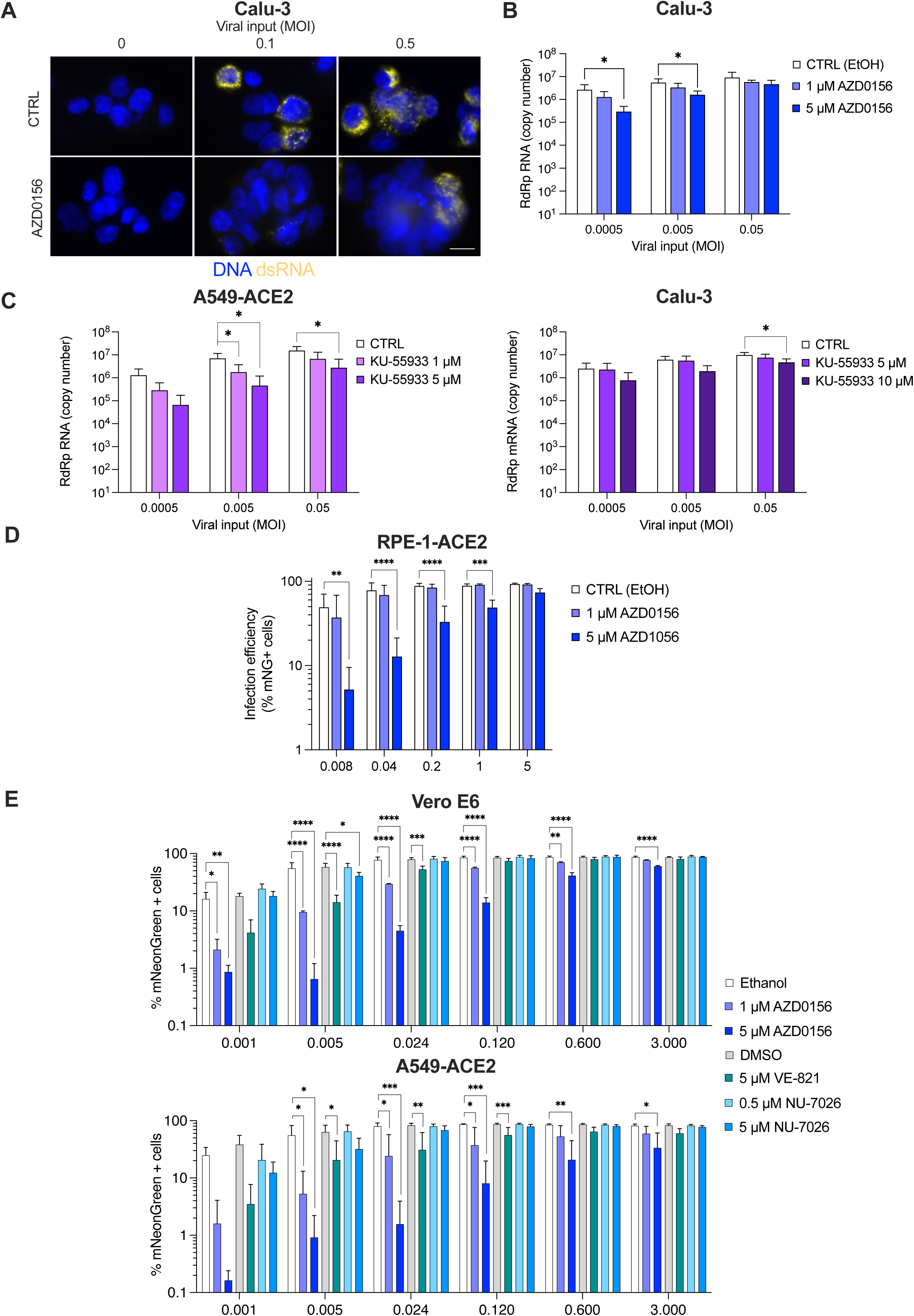
ATM inhibition inhibits SARS-CoV-2 replication. **A.** Calu-3 cells were incubated either with ethanol as a control or with ATM inhibitor AZD0156 at 5 μM 1 h prior to infection with SARS-CoV-2 at the indicated MOIs for 24 h before cell fixation and dsRNA was detected by immunofluorescence. Images are representative of 3 independent experiments. Images were acquired using a Zeiss AxioImager Z2 microscope. Scale bar is 15 µm. **B.** Viral RNA accumulation analysed by RT-qPCR (using RdRp specific primers and probe) in Calu-3 cells in the presence of the ATM inhibitor AZD0156 or not (ethanol control), as in Figure 3B. Mean and SD of biological quadruplicates. One-way ANOVA; *, *p* < 0.05. **C.** Viral RNA accumulation analysed by RT-qPCR (using RdRp specific primers and probe) in A549-ACE2 and in Calu-3 cells in the presence of the ATM inhibitor KU-55933 or not, as in Figure 3B. Mean and SD of 4 independent experiments. One-way ANOVA with Holm-Sidak’s test; *p* < 0.05; **, *p* < 0.01; ***, *p* < 0.001; ****, *p* < 0.0001. **D.** Impact of ATM inhibition on SARS-CoV-2(mNG) infection at the indicated MOIs in RPE-1-ACE2 cells. Infection efficiency was measured 2 days post-infection by flow cytometry. Mean and standard deviations of biological quadruplicates. Mean and SD of 3 independent 3 independent experiments. One-way ANOVA with Sidak’s test; *p* < 0.05; **, *p* < 0.01; ***, *p* < 0.001; ****, *p* < 0.0001. **E.** Vero E6 cells (top) and A549-ACE2 cells (bottom) were incubated for 1 h with the indicated drugs or controls (ethanol or DMSO) prior to infection with the indicated MOIs of SARS-CoV-2(mNG) and replication efficiency was monitored using flow cytometry 2 days later (AZD0156: ATM inhibitor; VE-821: ATR inhibitor; NU-7026: DNAPKc inhibitor). Mean and SD of 3 independent experiments. One-way ANOVA with Sidak’s test; *p* < 0.05; **, *p* < 0.01; ***, *p* < 0.001; ****, *p* < 0.0001 (when nothing is indicated, this is non-significant).

**Figure S4.**
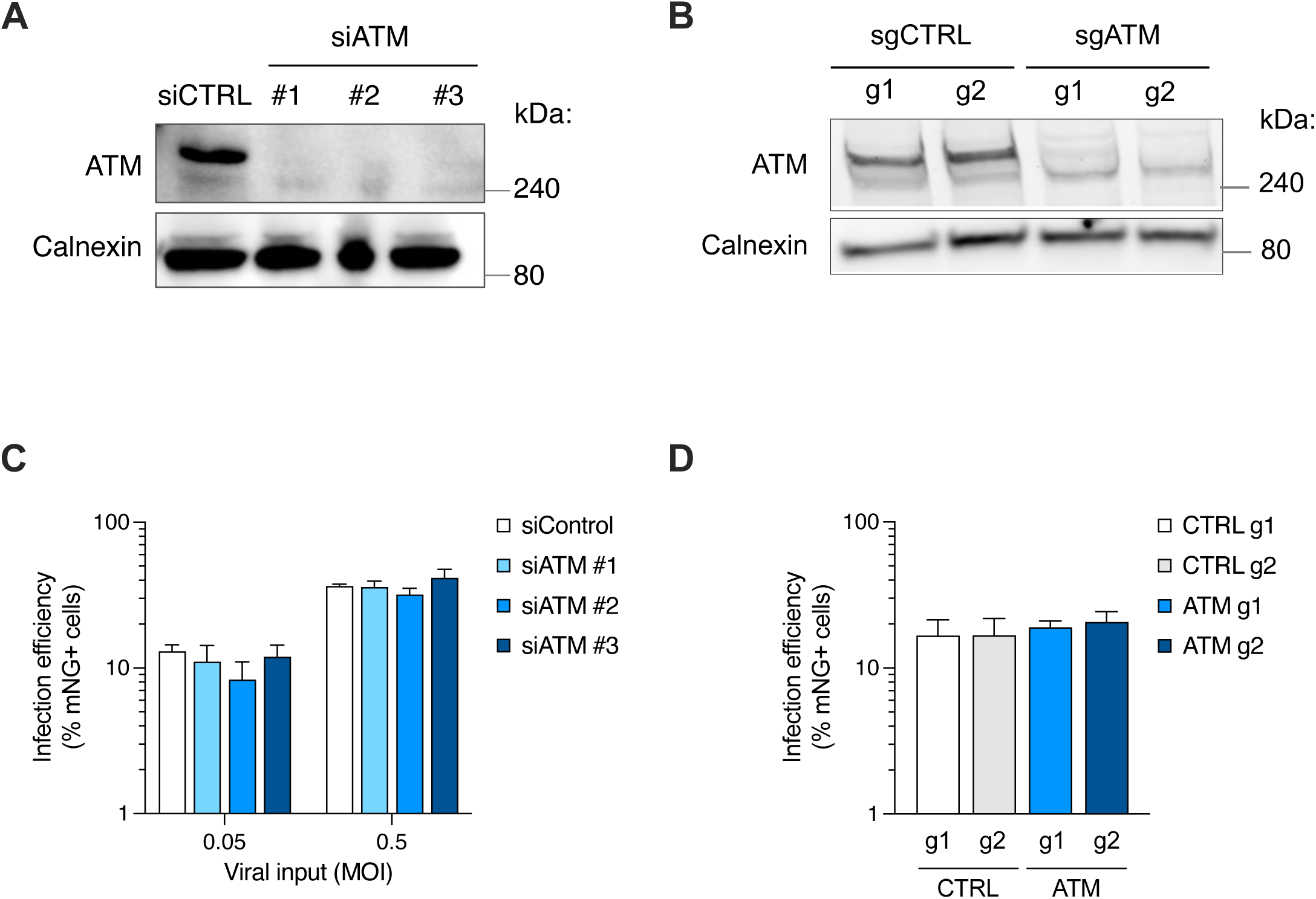
ATM depletion does not protect against SARS-CoV-2 replication in A549-ACE2 cells. **A.** Western blot showing siRNA-mediated ATM depletion 72 h post-transfection with three different siRNA targeting ATM (#1 to #3) as compared to siControl (siCTRL) in A549-ACE2 cells. Calnexin is used as a loading control. **B.** Western blot showing ATM expression in A549-ACE2 cells edited by CRISPR-Cas9 using two guides against *ATM* (sgATM), or two control guide RNAs (sgCTRL). **C.** Impact of ATM depletion by siRNA on SARS-CoV-2(mNG) infection at the indicated MOIs in A549-ACE2 cells. Infection efficiency was measured 2 days post-infection by flow cytometry. Mean and SD of 3 independent experiments. **D.** Impact of ATM depletion by CRISPR-Cas9 KO on SARS-CoV-2(mNG) infection at the MOI 0.2 in A549-ACE2 cells. Infection efficiency was measured 2 days post-infection by flow cytometry. Mean and SD of 4 independent experiments (performed in 2 independent series of KO cell populations).

## Notes

### Competing Interest Statement

The authors have declared no competing interest.

